# Optimizing Cost-Effective Gene Expression Phenotyping Approaches in Cattle Using 3′ mRNA Sequencing

**DOI:** 10.1101/2024.06.18.599599

**Authors:** Ruwaa I. Mohamed, Taylor B. Ault-Seay, Sonia Moisa, Jonathan Beever, Agustín G. Ríus, Troy Rowan

**Affiliations:** Genome Science and Technology Program, Bredesen Center, University of Tennessee, Knoxville, TN, USA; Animal Science Department, University of Tennessee Institute of Agriculture (UTIA), Knoxville, TN, USA

**Keywords:** Molecular phenotyping, gene expression, high-throughput, livestock, transcriptomics

## Abstract

**Background:** Genetic and genomic selection programs require large numbers of phenotypes observed for animals in shared environments. Direct measurements of phenotypes like meat quality, methane emission, and disease susceptibility are difficult and expensive to measure at scale but are critically important to livestock production. Our work leans on our understanding of the “Central Dogma” of molecular genetics to leverage molecular intermediates as cheaply-measured proxies of organism-level phenotypes. The rapidly declining cost of next-generation sequencing presents opportunities for population-level molecular phenotyping. While the cost of whole transcriptome sequencing has declined recently, its required sequencing depth still makes it an expensive choice for wide-scale molecular phenotyping. We aim to optimize 3′ mRNA sequencing (3′ mRNA-Seq) approaches for collecting cost-effective proxy molecular phenotypes for cattle from easy-to-collect tissue samples (*i.e.*, whole blood). We used matched 3′ mRNA-Seq samples for 15 Holstein male calves in a heat stress trail to identify the 1) best library preparation kit (Takara SMART-Seq v4 3′ DE and Lexogen QuantSeq) and 2) optimal sequencing depth (0.5 to 20 million reads/sample) to capture gene expression phenotypes most cost-effectively.

**Results:** Takara SMART-Seq v4 3′ DE outperformed Lexogen QuantSeq libraries across all metrics: number of quality reads, expressed genes, informative genes, differentially expressed genes, and 3′ biased intragenic variants. Serial downsampling analyses identified that as few as 8.0 million reads per sample could effectively capture most of the between-sample variation in gene expression. However, progressively more reads did provide marginal increases in recall across metrics. These 3′ mRNA-Seq reads can also capture animal genotypes that could be used as the basis for downstream imputation. The 10 million read downsampled groups called an average of 104,386 SNPs and 20,131 INDELs, many of which segregate at moderate minor allele frequencies in the population.

**Conclusion:** This work demonstrates that 3′ mRNA-Seq with Takara SMART-Seq v4 3′ DE can provide an incredibly cost-effective (<$25/sample) approach to quantifying molecular phenotypes (gene expression) while discovering sufficient variation for use in genotype imputation. Ongoing work is evaluating the accuracy of imputation and the ability of much larger datasets to predict individual animal phenotypes.

## Background

In livestock populations, selection decisions are made mainly based on statistical estimates of an animal’s genetic merit in the form of an estimated breeding value (EBV) [1]. Using EBV-based selection has led to remarkable genetic gain over relatively short periods. These genetic evaluations rely on large numbers of phenotypes measured on individuals in shared environments throughout the population [2], [3]. In most cases where genetic predictions do not exist for an economically-relevant trait, it is due to a lack of phenotypic measurements [4]. Some phenotypes, such as methane emission, disease susceptibility, or metabolic efficiency, are exceedingly challenging to quantify at the scale needed for genetic evaluation [5], [6], [7]. Future improvements to these economically important and sustainability-related traits will rely on novel approaches to collecting indicator phenotypes [8].

One possible solution to this challenge is to use molecular measurements (e.g., gene expression, protein, or metabolite abundance) as high-dimensional proxies for economically significant phenotypes [9]. These molecular measurements could be used in place of or alongside organismal phenotypes in genetic evaluations or in more complex models that predict animal phenotypes rather than additive genetic merit [10], [11], [12], [13]. Proxy phenotypes collected from milk samples via mid-infrared (MIR) spectroscopy are used as indicators for multiple efficiency and health traits in the dairy industry [14]. A similar proxy does not yet exist for beef cattle. As with any complex trait, genetic predictions that leverage intermediate phenotypes will also require large numbers of samples to be useful. As such, tissue collection must be accessible, and molecular data generation must be cheap. At present, the cost of whole transcriptome sequencing and proteomics remains prohibitively expensive. To equip future analyses with population-scale molecular phenotype data, we propose an optimized approach to quantifying gene expression via 3′ mRNA sequencing (3′ mRNA-Seq) in whole blood.

Whole transcript sequencing (mRNA-Seq) and 3′ mRNA-Seq are two related transcriptomic approaches, each offering unique representations of an individual’s gene expression landscape. While both approaches start by capturing mRNA of expressed genes, whole transcriptome sequencing libraries usually involve random fragmentation of entire genes, followed by fragment size selection for sequencing (Figure 1A). As a result, sequencing reads are distributed evenly across full transcripts. This can result in a biased overrepresentation of longer genes/transcripts amongst reads because they result in more fragments-per-transcript compared to shorter genes/transcripts [15]. This biased overrepresentation of longer genes can negatively affect the detection of differentially expressed genes (DEGs) and downstream analyses of biological functions like gene ontology (GO) enrichments and pathways analysis [16]. This requires that whole RNA sequencing analyses normalize the fragment counts per gene based on gene length and sequencing depth.

**Figure 1:**
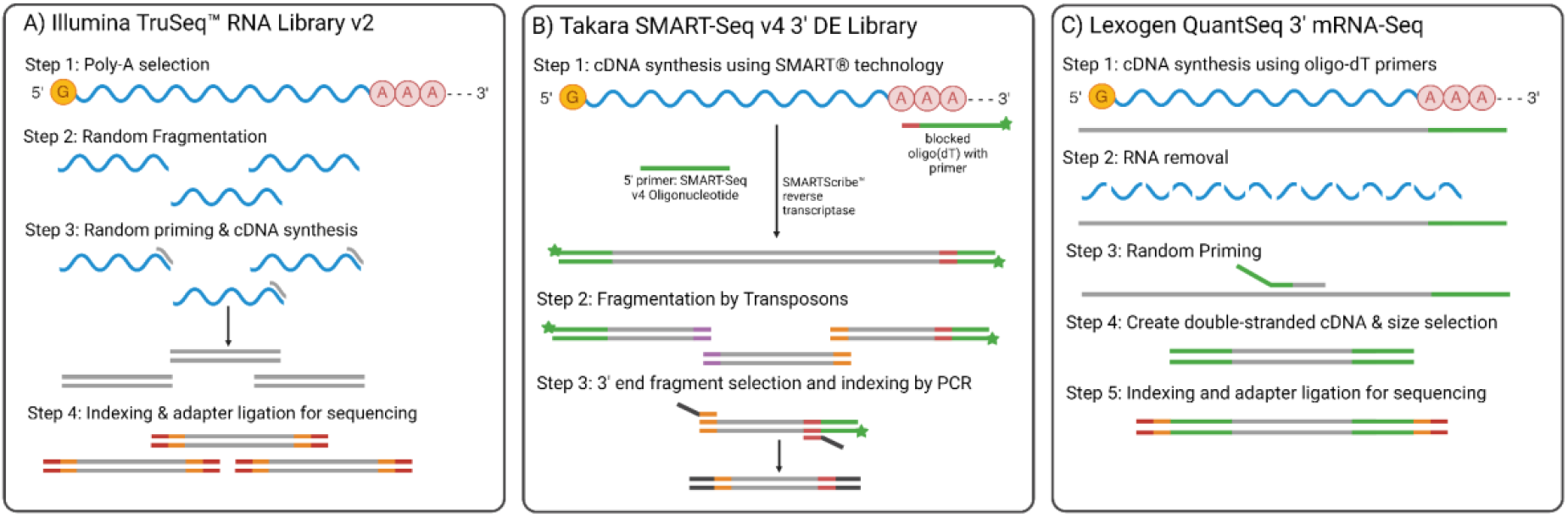
Comparison of the library preparation approaches for whole transcriptome sequencing and 3′ mRNA-Seq. A) Library preparation for Illumina TruSeq RNA library preparation used for whole transcriptome sequencing. B) Library preparation protocol for Takara SMART-Seq v4 DE approach used for 3′ biased sequencing. C) Library preparation protocol for Lexogen QuantSeq 3′ mRNA-Seq.

Comparatively, 3′ mRNA-Seq approaches do not fragment the captured transcripts and sequence multiple fragments from the same transcript. Instead, 3′ mRNA-Seq involves a biased library preparation approach that generates reads from the 3′ end of mRNA molecules and creates a cDNA library from only the fragment containing the poly-A tail (Figure 1B,C). This results in an unbiased representation of long and short transcripts in the sequencing library [15]. As a result, gene expression analysis of 3′ mRNA-Seq does not require normalization based on gene length. The most important benefit of 3′ mRNA-Seq is that gene expression can be quantified with a fraction of the reads necessary in a full transcriptome study due to a reduction in the number of redundant reads that belong to the same transcript [15]. The main drawback of 3′ mRNA-Seq is that reads cannot be used to identify novel transcripts or alternative splicing events. Further, it can only call 3′ biased intragenic variants in cases where genomic variants are of interest. Multiple commercial library preparation approaches are available for 3′ mRNA-Seq, including Tag-seq [17], [18], MustSeq [19], QuantSeq (Lexogen) [20], and SMART-Seq v4 3′ DE (Takara). The Lexogen QuantSeq library is one of the most widely used 3′ mRNA-Seq libraries across studies, used in over 1,600 publications in the past decade [21].

The literature is conflicting on whether 3′ mRNA-Seq or whole transcriptome sequencing approaches are better equipped to detect differentially expressed genes (DEGs) [15]. Some side-by-side comparisons found that full transcriptomes were more effective at detecting DEGs [15], while others have identified 3′ mRNA-Seq as the superior approach [19], [20], [22], [23]. However, the majority of comparisons suggest that the libraries are equally capable of detecting both DEGs and expressed genes [17], [18], [24], [25], [26], [27]. Finally, compared with whole transcriptome sequencing, 3′ mRNA-Seq represents the major cost savings needed for widespread molecular phenotyping applications [15], [18].

This project aimed to identify best practices for carrying out cost-effective 3′ mRNA-Seq for eventual application in the molecular phenotyping of livestock. We explored how different library preparation approaches and sequencing depths affect the quality and amount of information generated. Here, we compare two popular library preparation approaches for 3′ mRNA-Seq: Lexogen QuantSeq and Takara SMART-Seq v4 3′ DE. We also use an iterative downsampling approach to simulate the impacts of different sequencing depths. This allows us to identify the minimum depth needed to capture the optimal amount of variation in transcript quantities. Finally, we explore our ability to call variants from 3′ biased sequencing reads. The ability to impute genotypes based on this reduced-representation gene expression data would be invaluable for many genotype-to-phenotype applications in plants and livestock. Coupled with developments in laboratory automation, the best practices identified by this work could make molecular phenotyping practical for livestock genetic evaluations.

## Methods

### Sample collection and RNA extraction

We opportunistically collected whole blood samples from 15 Holstein male calves housed under climate control rooms [29] in the East Tennessee Research and Education Center – JRTU undergoing an acute heat stress trial for an unrelated project (Yu et al., under revision). We handled all animals according to the University of Tennessee’s Institutional Animal Care and Use Committee Protocol 2851-0921. Fifteen whole blood samples were collected from the cattle at 6:30 before heat exposure, and 14 samples were collected at 18:30 immediately after 12 hours of heat exposure (Yu et al., under revision). Climate in the room was obtained following our previous work in heat-stressed claves [29]. 10 mL of blood was mixed with 30 mL of 1X NH_4_Cl red blood cell lysis buffer and was centrifuged at 2000 *×g* for 10 minutes. The supernatant was aspirated and cell pellets resuspended in 1.2 mL of Trizol. RNA was isolated according to the protocol detailed in Rio *et al.* [28]. RNA purification and genomic DNA removal were performed using the Zymo RNA Clean and Concentrator kit according to manufacturer protocol.

### Library Preparation and Sequencing

Sequencing libraries were prepared from isolated RNA using two different kits: Takara SMART-Seq v4 3′ DE (Takara) and Lexogen QuantSeq 3′ (Lexogen) (Supplementary Table S1) as per the manufacturer’s instructions. Final library quality and concentrations for pooling were evaluated using the Agilent Tapestation 4200 system. For each kit, libraries for all 27 samples were pooled to achieve equal concentrations (Supplementary table 1). For the Lexogen pool, we targeted adding 5 ng of each sample to the pool. For Takara, we targeted adding 3.2 ng of each sample to the pool. They were sequenced on a single SP flow cell on the Illumina Novaseq6000 (University of Tennessee Genomics Core - Knoxville, TN) with a 200-cycle v1.5 reagent kit. For Takara 3′ libraries, Read 1 was 150 bp, and Read 2 consisted of 26 bp for demultiplexing. Lexogen 3′ libraries were sequenced as single-end with 150 bp reads.

### Sequence Processing & Gene Expression Quantification

Only forward reads were used for sequence analysis, as reverse reads contained only indices for demultiplexing for the Takara kit. For all samples, Trimmomatic (v.0.39) was used for trimming and filtering with the following parameters: (LEADING:5 TRAILING:5 SLIDINGWINDOW:5:20 MINLEN:30) to trim bases from each end of each read that has a quality ≤ 5, trim at the first occurrence of sliding window of size 5 with average quality ≤ 20, and filter out the reads that are shorter than 30 bp after the trimming [30]. The quality of reads before and after trimming & filtering was visualized and evaluated using FastQC (v.0.11.9) [31] and MultiQC (v.1.14) [32] tools. The STAR alignment software (v.2.7.10b) [33] was used to index the *Bos taurus* genome (ARS-UCD2.0; Jul 2023) [34] - which was obtained from NCBI - with sjdbOverhang of 149 bp. STAR was then used to map filtered reads to the indexed genome. The quantMode “GeneCounts” was used in STAR to quantify the number of reads mapped to the annotated genes.

### Differential Gene Expression Analysis & Functional Enrichment Analysis

We evaluated the sensitivity of each test condition (library preparation method and downsampled read number) to detect expressed genes and differentially expressed genes. The number of expressed genes (count > 0) and informative genes (count > 10 in 50% of the samples) were calculated in R (R 4.2.1 "Funny-Looking Kid") using the GeneCounts generated by the STAR tool. Differentially Expressed Genes (DEGs) were identified using the DESeq2 R package (v.1.40.2) with an FDR-corrected alpha threshold of 0.05 [35]. No log fold change threshold was used for DEG identification. DEGs were plotted using the Bioconductor EnhancedVolcano library [36].

### Variant Calling

We utilized the GATK best practices for calling variants from RNA-sequencing data to identify genomic variation in areas covered by 3′ mRNA-Seq reads [37], [38]. STAR was used to remap the reads with a per-sample 2-pass mapping step. Samples from the same animal (from before and after heat exposure) were merged using SAMtools (v.1.20) [39]. We used the GATK tool (v.4.3.0.0) [38] and picard tools (v.2.27.4-0) [40] to: 1) Assign all reads from each animal to a single read group identified by the animal ID using the AddOrReplaceReadGroup tool, 2) mark duplicated reads in each of the merged samples, 3) split the reads that contain Ns in their cigar string using SplitNCigarReads tool, 4) Call variants within each of the merged samples using HaplotypeCaller with a confidence threshold of 20, 5) combine the variants from all samples across all chromosomes and known scaffolds in the genome using GenomicsDBImport, 6) perform joint variant calling from all samples using GenotypeGVCFs, 7) Separate SNPs and INDELs using SelectVariants, and 8) Hard-filter SNPs and INDELs, separately, with VariantFiltration tool with the following parameters for SNPs (QD<2.0, FS > 60.0, MQ < 40.0, MQRandSum < -12.5, ReadPosRankSum < -8.0, and SOR > 3.0) and the following parameters for INDELs (QD<2.0, FS > 200.0, QUAL < 30.0, and ReadPosRankSum < -20.0). Sites present in less than or equal to one animal were removed from the dataset.

### Downsampling for sequencing-depth benchmarking

We used the Seqtk tool (v.1.4) [41] to generate ten random replicates of downsampled read sets simulating seven ascending sequencing depths from raw FASTQ files for each sample. We simulated this downsampling using ten random seeds (127, 2, 5, 7, 9, 11, 12, 81, 21, 47) at each of seven different sequencing depths: 0.5M, 1M, 2M, 5M, 7.5M, 10M, and 12M reads/sample. Higher sequencing depth downsampling past 12 M reads was not possible for all samples. We performed the same analysis described above (alignment, gene expression counting, DEG analysis, and variant calling) to quantify the relative performance of each sequencing depth across replicates.

The results of the downsampling were quantified using precision, recall, and F-score where the whole dataset’s results were considered the true positive set. Precision is the measure of the positive predicted value (PPV), recall is the measure of sensitivity, and F-score is the harmonic mean of precision and recall [42]. The three metrics were calculated using the following equations:

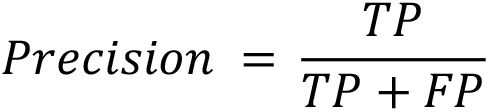

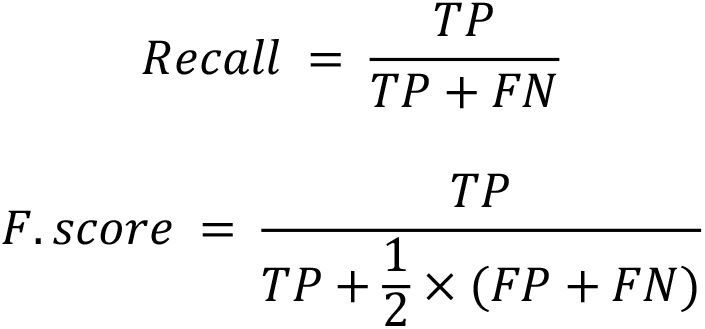

Where

- True positive (TP) is the number of genes present in both the subsample and the full dataset.

- False positive (FP) is the number of genes present only in the subsample.

- False negative (FN) is the number of genes present only in the full dataset but not the subsample.

The plateau value, indicating the point where additional reads failed to uncover additional genes for each metric, was estimated by fitting the downsampling results from all ten replicates to a Self-Starting Nls Asymptotic Regression Model (SSasymp function from the stats package in R) with the following equation: 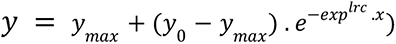, Where

- *y_max_* is the asymptote value on the y-axis,

- *y*_0_ is the response when *x* = 0, and

- *lrc* is the natural logarithm of the rate constant.

The plateau’s initial point (*i.e.,* saturation sequencing depth) was calculated by setting the derivative of this model to 0.0001, which represents an increase of one gene from adding 10,000 extra raw reads per sample (*i.e.,* marginal information gain).

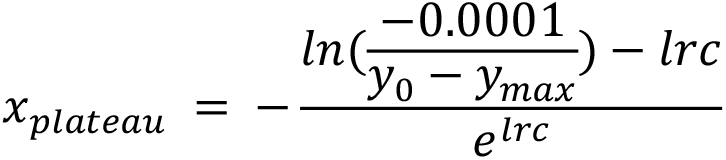

## Results

### Comparing Takara and Lexogen libraries

Sequencing depth was similar for both library preparation kits (p-value 0.9761; Welch two-sample t-test) with a median of 18.6 million (range: 12.5 M - 35.6 M) and 16.4 million (range: 7.3 M - 95.7 M) raw reads per sample for the Takara and Lexogen libraries, respectively (Supplementary Figure S1). One sample sequenced with the Lexogen library had significantly more reads than all other samples (>5 standard deviations from the mean) because it was inadvertently loaded to a higher concentration. On average, quality control filters dropped significantly fewer reads for the Takara library (1.94% of the raw reads) compared with the Lexogen library (2.39% of the raw reads) (p-value < 0.0001; Welch two-sample t-test) (Figure 2A). When mapped to the cattle reference genome, the reads from Lexogen libraries had a significantly higher mapping rate on average, with 87.95% of the raw reads where uniquely mapped compared to Takara libraries with 75.56% uniquely mapped reads (p-value < 0.0001; Welch two-sample t-test) (Figure 2B). This was driven by more reads in the Takara data being unmapped because they were too short (not shown). More stringent filtering criteria would decrease the rate of unmapped reads from both sequencing libraries.

**Figure 2:**
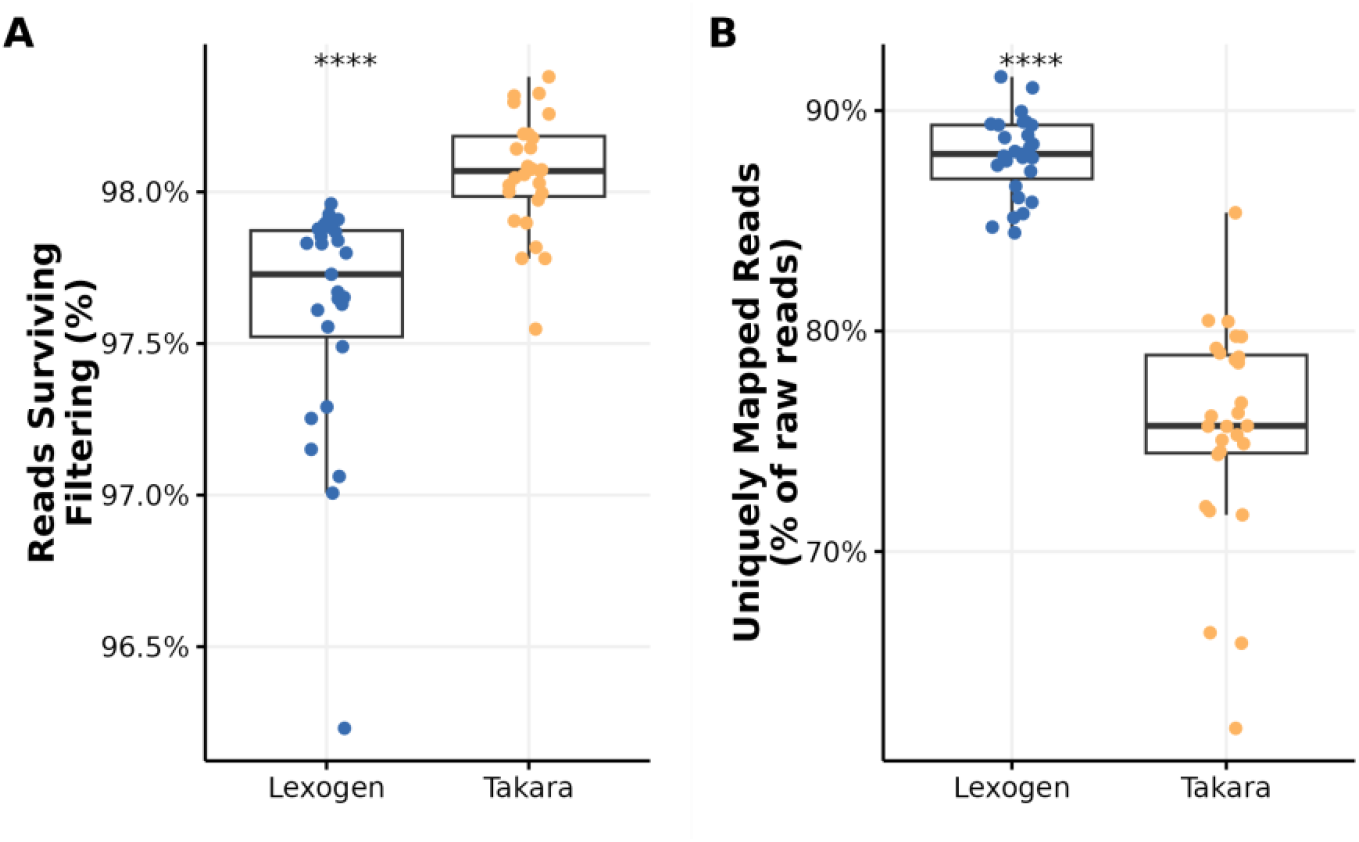
Comparison of Takara and Lexogen filtering outcomes and mapping rates. (A) Boxplots represent the percentage of the raw reads that survived filtering from both sequencing libraries. (B) Boxplots representing the percentage of the raw reads that survived filtering and were uniquely mapped to the cattle reference genome. (****: p <= 0.0001)

Despite Lexogen libraries having a higher mapping rate, Takara libraries captured significantly more expressed genes than Lexogen libraries (p-value < 0.0001; Welch two-sample t-test). We considered “expressed genes” as genes to which at least a single read aligned. Takara libraries detected an average of 16,957 expressed genes per sample (minimum = 15,921; maximum = 18,532), while Lexogen libraries averaged only 13,417 (minimum = 9,273; maximum =16,033) (Figure 3A). Using both libraries, the number of expressed genes from all animals (Figure 3A black stars) is greater than the number of expressed genes from any individual, with a total of 24,607 expressed genes from Takara libraries and 21,730 expressed genes from Lexogon libraries. Our definition of an expressed gene was quite liberal, meaning that many were captured only in one or a handful of individuals, which could mean they would not be as informative as molecular phenotypes (Figure 3C, D). We defined a more conservative set of “informative genes” as genes where at least ten reads mapped to the gene in at least 50% of the samples. Takara libraries captured significantly more informative genes per sample (mean = 11,824) than Lexogen libraries (mean = 9,904) (p-value < 0.0001; Welch two-sample t-test) (Figure 3B). Takara libraries captured 12,351 informative genes from all samples, whereas Lexogen libraries captured only 10,997 informative genes (Figure 3B black stars).

**Figure 3:**
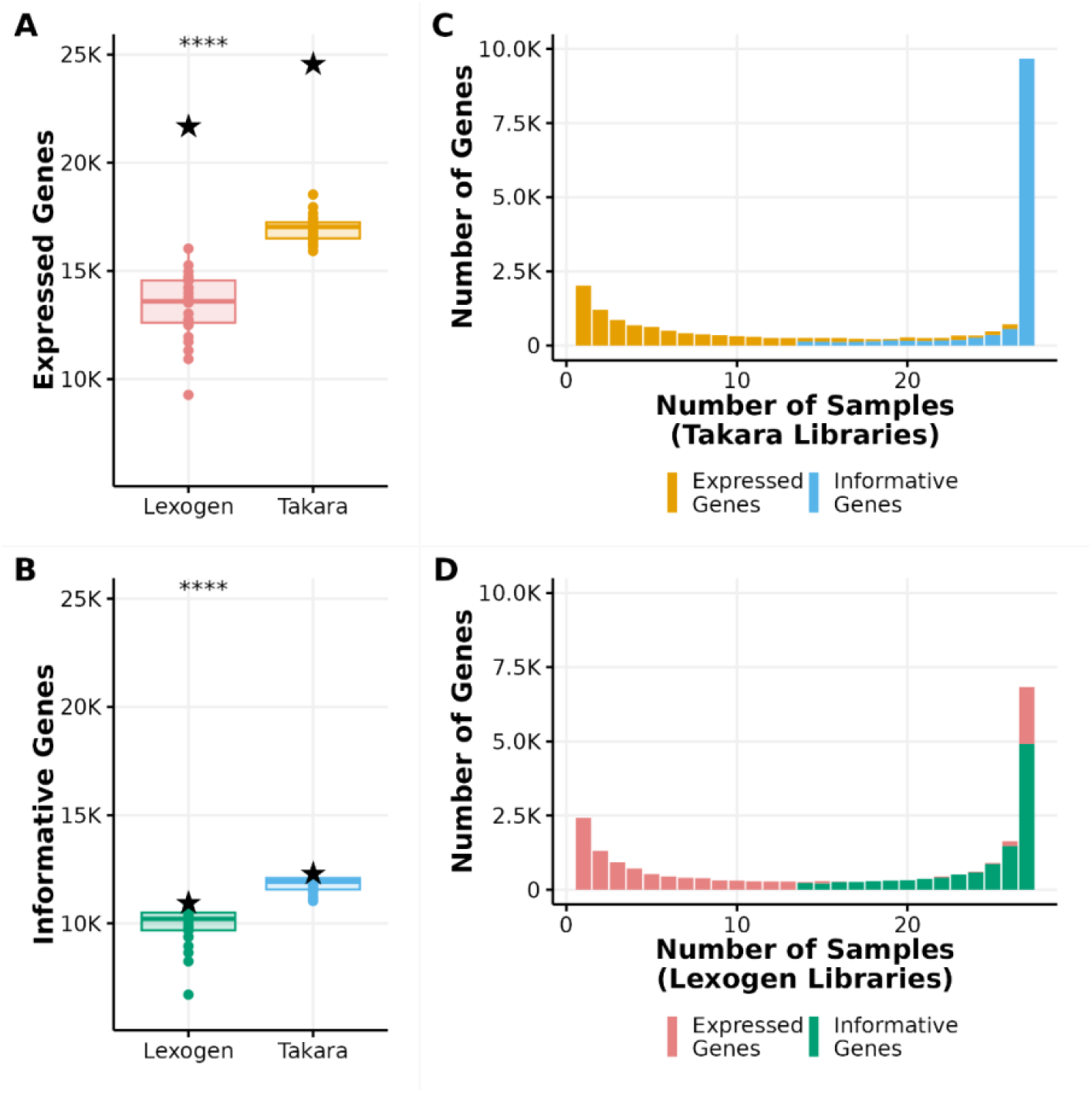
Comparison of expressed and informative genes detected by Takara and Lexogen library preparation kits. (A) Boxplots representing the numbers of expressed genes (genes with at least one read mapped) for Takara and Lexogen libraries. Each dot represents a sample, and the black star represents the number of expressed genes identified in the full dataset for each library preparation method. (B) Boxplots representing the number of informative genes (genes with at least ten mapped reads in 50% of the samples) for Takara and Lexogen libraries. Each dot represents a sample, and the black star represents the number of informative genes present in at least 50% of the samples identified in the full dataset. (C, D) Distribution of the number of expressed genes and informative genes captured by <u>Takara libraries</u> (C) and <u>Lexogen libraries</u> (D) as a function of the number of individuals in which these genes were identified. (****: p <= 0.0001). The colors in subfigures (A) and (B) match the colors in subfigures (C) and (D), respectively.

When we modeled the heat stress response between samples, Takara enabled us to identify more differentially expressed genes (DEGs) than Lexogen. Takara libraries identified 4,821 DEGs (3,142 up-regulated genes and 1,679 down-regulated genes), whereas Lexogen libraries captured only 1,285 DEGs (1,025 up-regulated genes and 260 down-regulated genes) (Figure 4). Only 1,095 DEGs were consistently detected using both libraries (964 up-regulated genes and 131 down-regulated). Most of these DEGs had a small effect size as only 179 of the up-regulated genes had |*LFC*| ≥ 1 from the Takara libraries and only 47 up-regulated genes and 2 down-regulated genes from Lexogen libraries had |*LFC*| ≥ 1 (not shown). 94% of the upregulated genes and 50% of the downregulated genes captured by Lexogen were also captured by Takara libraries (Figure 4C).

**Figure 4:**
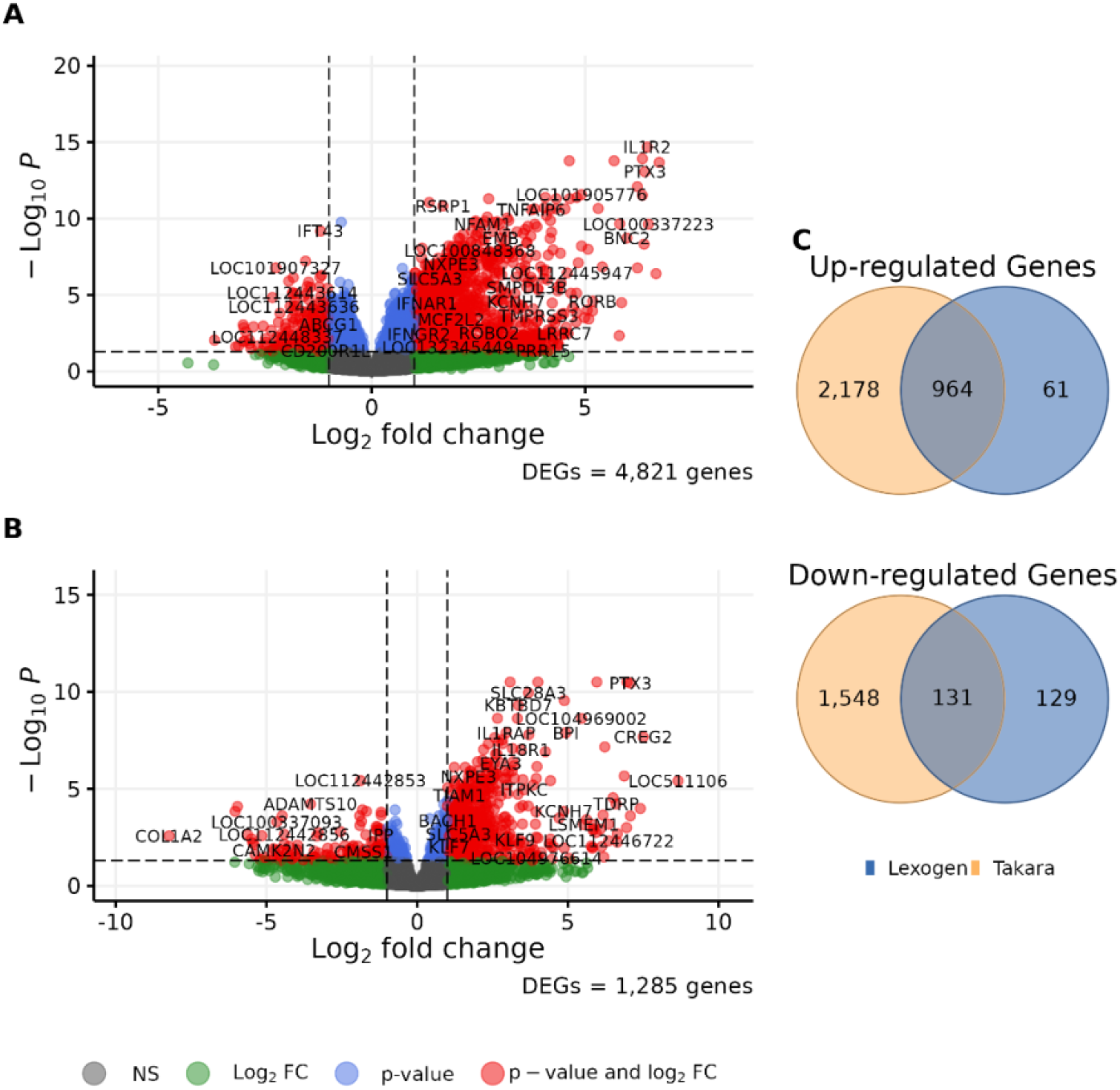
Volcano plots representing the differentially expressed genes of heat-stressed calves using A) Takara libraries and B) Lexogen libraries. The horizontal dashed line represents the adjusted p-value threshold of 0.05; all genes above the line are statistically significant. The vertical dashed lines represent |*LFC*| ≥ 1; all genes beyond the vertical lines have moderate-to-high effect size as a response to heat stress. Differentially expressed genes (red dots) are both statically significant and have moderate-to-high effect sizes. C) Number of DEGs detected by both library kits.

We called more genetic variants using reads from the Takara libraries than from Lexogen libraries. However, the quality of the called variants was not different between both library preparations, with a mean filtration survival rate of 0.76 from both libraries. After variant filtration and genotype filtration, we obtained 154,244 SNP sites and 29,711 INDEL sites from Takara library sequences, compared with 116,984 SNP sites and 16,407 INDEL sites from Lexogen libraries (Figure 5A). For Takara libraries, 137,570 SNP sites (89.19% of the called Takara SNP sites) were intragenic (overlapping with 13,322 annotated genes), with the majority of the SNP sites overlapping with informative genes (Figure 5A). Compared with 109,366 SNP sites (93.49% of the called Lexogen SNP sites) inragenic SNP sites (overlapping with 9,726 genes) from Lexogen libraries with the majority of the SNP sites overlapping with informative genes (Figure 5A). Interestingly, most of the called variant sites were unique to only one of the sequencing libraries, as only 32,245 SNP sites & 5,973 INDEL sites were called using both sequencing libraries (Figure 5B). Figure 5C shows the distribution of SNP sites called using both sequencing libraries. From both libraries, a higher density of SNP sites was called near the 3′ end of the gene body as expected (Figure 5D).

**Figure 5:**
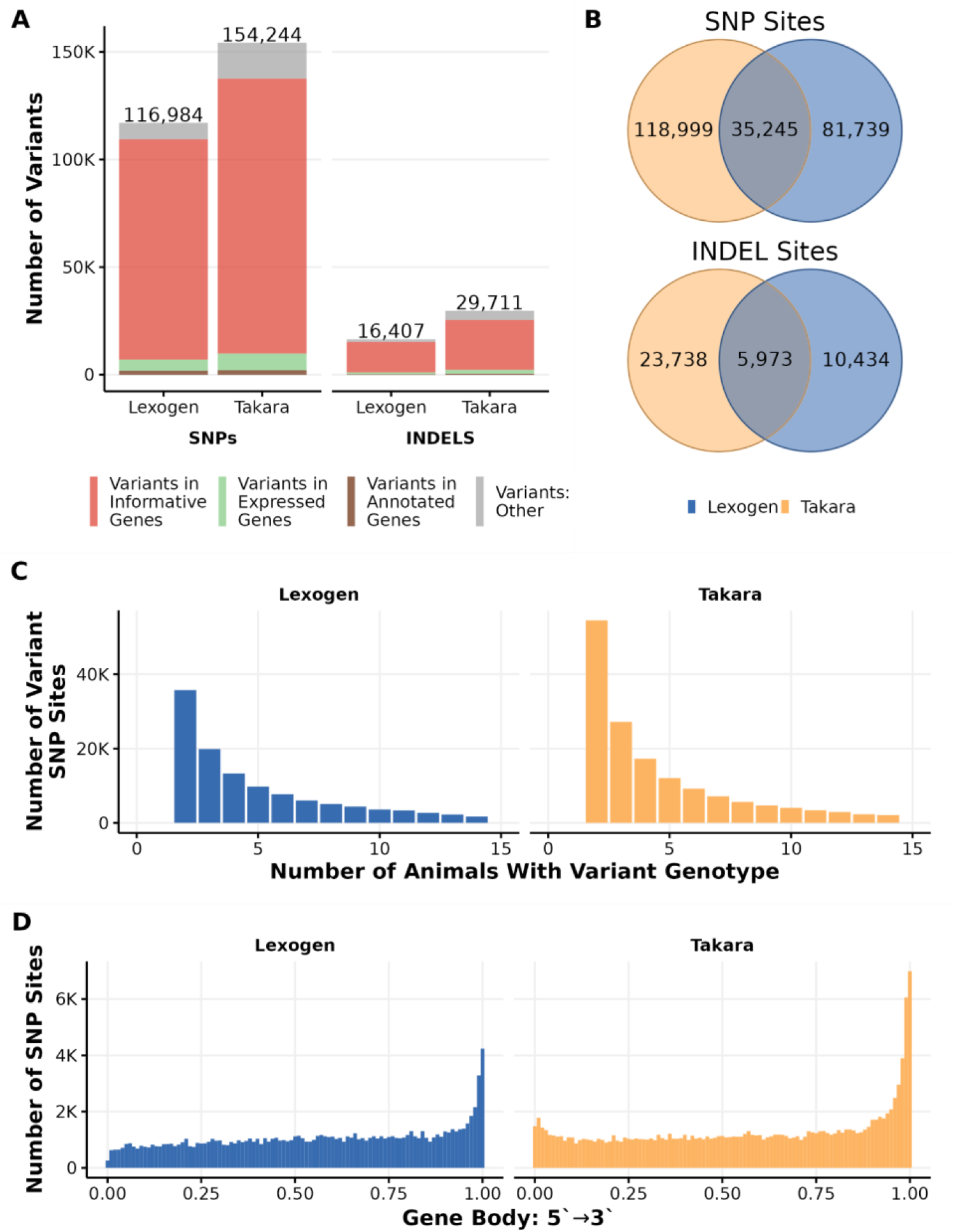
Variant calling results from Takara and Lexogen libraries. A) Bar chart depicting differences in the number and location of variant sites. B) The Venn diagram shows the number of shared and unique variant sites between both sequencing libraries. C) Distribution of the animals’ genotypes for the SNP sites. E) distribution of the called intragenic SNP sites across the gene body from 5′ → 3′ of the gene.

### Identifying an optimal read depth for 3′ mRNA-Seq in molecular phenotyping applications

After identifying Takara as the superior library preparation approach, we performed a series of read number-downsampling iterations to determine the optimal sequencing depth that balanced capturing expression variability while minimizing costs. The unique mapping rate and filtering rate were the same across all downsampling iterations, as it is driven by sample quality rather than read number. (Supplementary Figure S2 A, B). However, the number of expressed and informative genes per sample increased with sequencing depth (Supplementary Figure S2 C, D). The number of expressed genes and informative genes captured by all replicates increased exponentially as the sequencing depth increased (Figure 6 A, B). For expressed genes, asymptotic regression models plateau at 7,993,318 reads per sample, capturing 23,605 predicted expressed genes (turquoise dot in Figure 6A). For informative genes, the model starts to plateau at 8,357,463 reads per sample, capturing 11,137 predicted informative genes (turquoise dot in Figure 6B). The model begins to plateau at 6,565,915 reads per sample for DEGs, capturing 4,297 DEGs (turquoise dot in Figure 6C) out of 4,821 DEGs captured by the whole dataset. The model predicted that using the full number of reads generated (12,601,460) would only capture an additional 235 DEGs.

**Figure 6:**
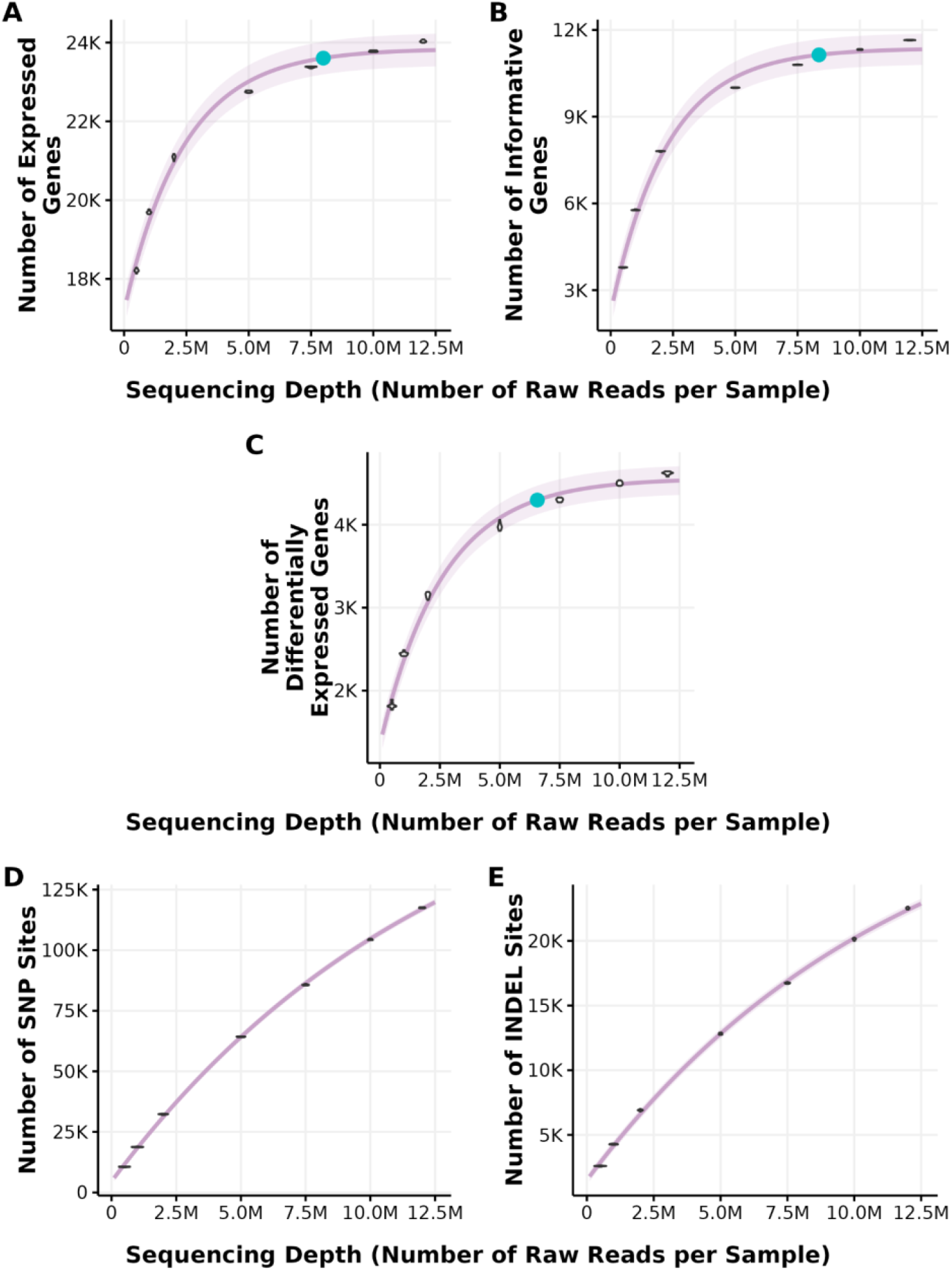
The effect of sequencing depth on capturing expressed genes, informative genes, DEGs, and variants. Each sequencing depth downsample consisted of ten repetitions (a cluster of 10 points from the downsampling subsets). Lilac lines are non-linear asymptotic regression models; each parameter was fit to a regression model independently. Turquoise dots represent the start of the plateau, defined by a slope of 0.0001, representing a gain of 1 gene/variant when increasing the sequencing depth by 10,000 reads per sample. (A) Number of expressed genes as a function of sequencing depth per sample. (B) Number of informative genes (expressed genes with at least 10 reads mapped to in at least 50% of the samples) as a function of sequencing depth per sample. (C) Number of differentially expressed genes (DEGs) as a function of sequencing depth per sample. (D) Number of SNP sites called in at least 2 but not all animals as a function of sequencing depth per sample. (E) Number of INDEL sites called in at least 2 but not all animals as a function of sequencing depth per sample.

The depth of sequencing also affected the number of variants that we were able to call from 3′ mRNA-Seq reads. Unlike gene expression, the number of filtered variants did not plateau at any sequencing depth, including at the full 12.5M reads per sample (Figure 6 D, E). We could not call variants at higher sequencing depth as only nine samples had a sequencing depth ≥ 20M, and only two samples had ≥ 30M reads. As expected, our analysis showed that the number of samples is as important as the number of reads in accurately calling variants. Using our limited dataset, the asymptotic regression model extrapolates the plateau to be around 64,396,412 reads per sample for calling SNP sites and 39,010,649 reads per sample for calling INDEL sites; however, extrapolation of regression models beyond the range of observed data should be approached with caution, as the reliability of predictions in these regions is often questionable. Using the optimal sequencing depth identified for identifying expressed and informative genes 8 million reads per sample), we called 90,343 SNP sites and 17,625 INDEL in the dataset. This represented 58.6% and 59.2% of total SNP and INDEL sites called with the complete dataset, respectively.

## Discussion

Our study demonstrates the potential of using 3′ mRNA sequencing (3′ mRNA-Seq) as a cost-effective method for molecular phenotyping in cattle. We primarily focus on identifying superior library preparation kits and optimizing sequencing depth to balance information content and cost. Our findings indicate that the Takara SMART-Seq v4 3′ DE library preparation kit outperforms the Lexogen QuantSeq kit across various metrics, including the number of quality reads, expressed genes detected, informative genes detected, differentially expressed genes identified, and intragenic variants called. This 3′-biased approach to sequencing can both increase sample sizes in differential gene expression analysis & eQTL studies and enable population-level molecular phenotyping. We anticipate that gene expression phenotypes can be used as high-dimensional indicators for other hard-to-measure traits in cattle, such as methane emissions, metabolic efficiency, reproductive predisposition, or disease susceptibility.

We primarily evaluated library preparation approaches and optimal sequencing depth by the number of informative genes detected. We defined informative genes as those with at least ten mapped counts that were detected in at least 50% of the samples. While detecting all expressed genes is important, this core set of genes will be necessary for extrapolating latent phenotypes across the wider population. We found that Takara libraries captured a significantly greater number of informative genes compared to Lexogen libraries. In addition to the comparative performance of the library preparation kits, our analysis of the sequencing depth revealed that the Takara SMART-Seq v4 3′ DE library preparation kit’s sensitivity to capture expressed genes and informative genes from cattle whole blood saturates at a sequencing depth of around 8 million reads per sample. We observed that increasing the read depth beyond 8 million reads provided only marginal gains in the recall of gene expression metrics, suggesting a point of diminishing returns in terms of cost versus data richness. In contrast, *Xiong et al.* found that the sensitivity of the 3′ mRNA-Seq library to detect expressed genes saturates at 2-3 million reads per sample from human primary cardiomyocyte cell lines [24]. That said, the optimal sequencing depth is likely species- and tissue-specific, reflecting transcription activity in different types of cells and organisms.

The ability to call variants from 3′ mRNA-Seq reads also demonstrated a strong dependence on read depth and number of samples. Our results in a small set of animals showed that while gene expression metrics plateaued at lower read depths, variant identification continued to benefit from increased sequencing depth without reaching an apparent plateau within the range tested. Further sequencing would likely result in greater proportions of gene bodies to be sequenced, resulting in more variants. That said, the 100,000+ high-quality variants called in the 3′ portion of genes are likely sufficient to enable genotype imputation when adequately constructed reference panels are available.

Our study highlights the importance of carefully optimizing sequencing protocols to balance cost and data quality. Using 3′ mRNA-Seq with the Takara SMART-Seq v4 3′ DE kit offers a promising approach for large-scale molecular phenotyping in livestock, providing sufficient data quality at a substantially lower cost than whole transcriptome sequencing. Using fractional quantities of reagents for library preparations could provide even further reductions in cost. This approach enables the capture of large-scale gene expression data and may also facilitate genotype imputation, thereby enhancing the utility of molecular data in genetic evaluations and breeding programs.

## Conclusion

In conclusion, we show that 3′ mRNA-Seq is a cost-efficient (<$25/sample) approach to studying and representing complex traits in cattle through phenotyping by gene expression. In our samples, the Takara SMART-seq v4 library was superior to the Lexogen QuantSeq library in capturing expressed, informative, and differentially expressed genes, as well as calling sequence variants. We have also shown that 8 million reads per sample effectively capture most of the inter-sample variation in gene expression with a marginal increase in the number of expressed genes with increasing read depth. However, ongoing work is identifying the optimal sequencing depth for variant calling from 3′ mRNA-Seq and imputation.

## Funding

This work was funded in part by an AG2PI Round 2 Seed Grant. The AG2PI project is supported by USDA-NIFA awards 2020-70412-32615 and 2021-70412-35233.

## List of abbreviations

DEGs: Differentially Expressed Genes
FP: False Positive
FN: False Negative
GATK: Genome Analysis Toolkit
GO: Gene Ontology
INDEL: Insertions/Deletions
MAF: Minor Allele Frequency
SNP: Single Nucleotide Polymorphism
TP: True Positive

---

**Figure S1:**
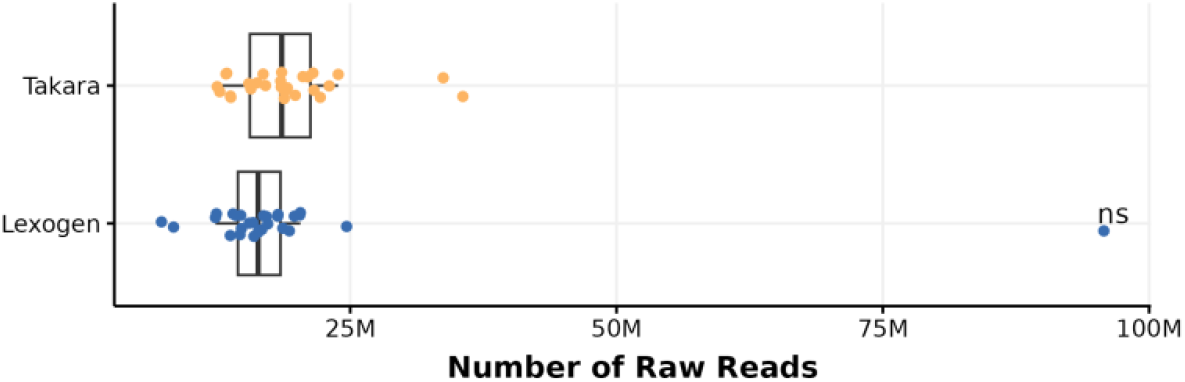
Number of raw reads obtained from sequencing with Takara and Lexogen 3′ mRNA-Seq library kits (p-value = 0.9761). We obtained a median of 18.6 M (range: 12.5 M - 35.6 M) from Takara libraries (top) and 16.4 M (range: 7.3 M - 95.7 M) raw reads per sample from Lexogen libraries (bottom).

**Figure S2:**
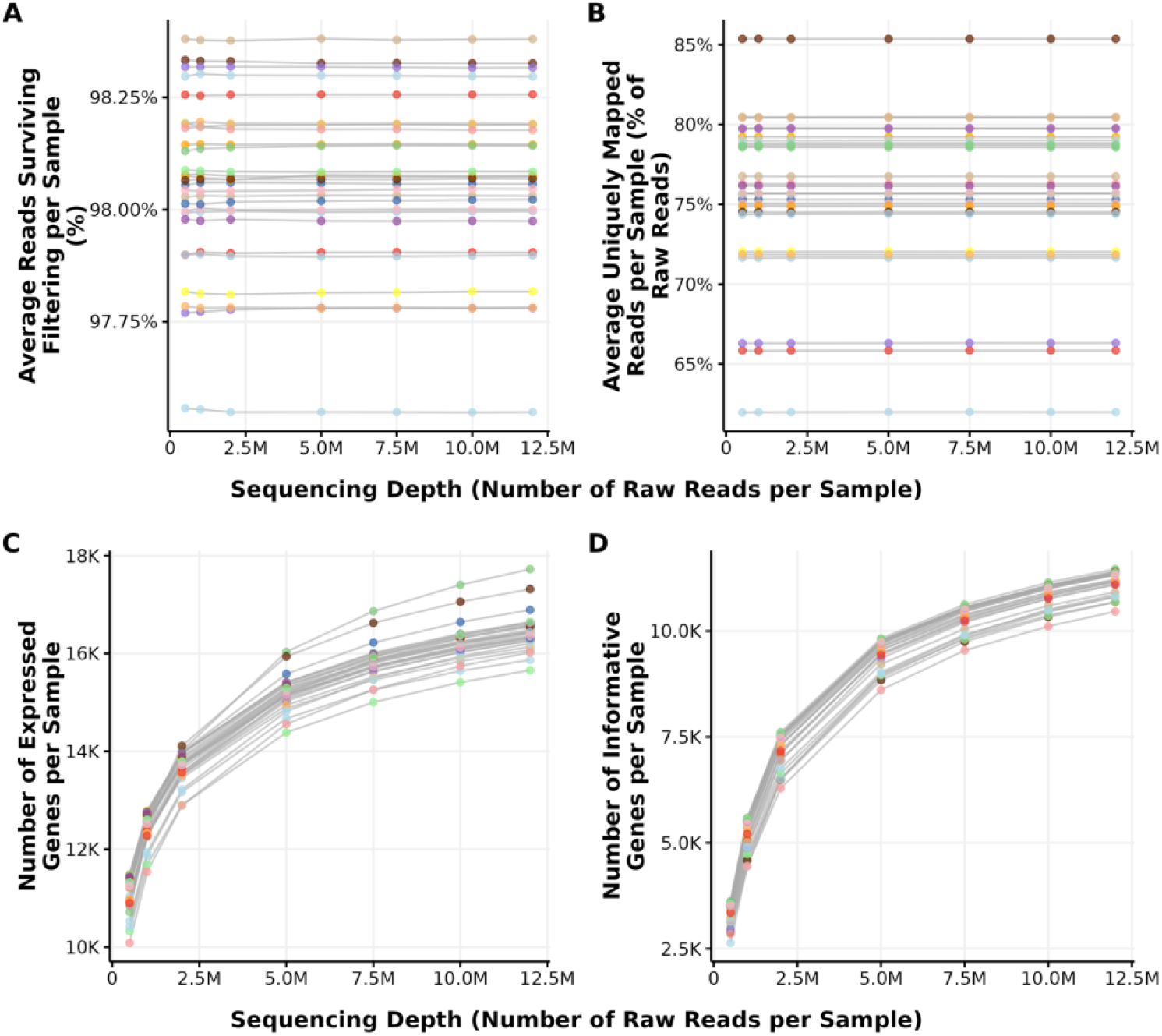
Effect of sequencing depth for Takara libraries on per-sample metrics. Each line represents a sample, and each dot is the average of ten replicates. Each color represents one animal. (A) Average read filtration rate (%) as a function of the number of raw reads per sample. (B) Per sample average uniquely mapped read (% of raw reads) as a function of the number of raw reads per sample. Both the filtering rate and mapping rate are entirely dependent on the sample and the quality of the sample (handling, extraction protocol, library preparation, sequencer, etc.) and independent of the sequencing depth. (C. D) Number of expressed genes (C) and informative genes (D) per sample as a function of sequencing depth (Number of raw reads per sample). Both metrics are dependent on the sequencing depth, similar to Figure 6.

**Figure S3:**
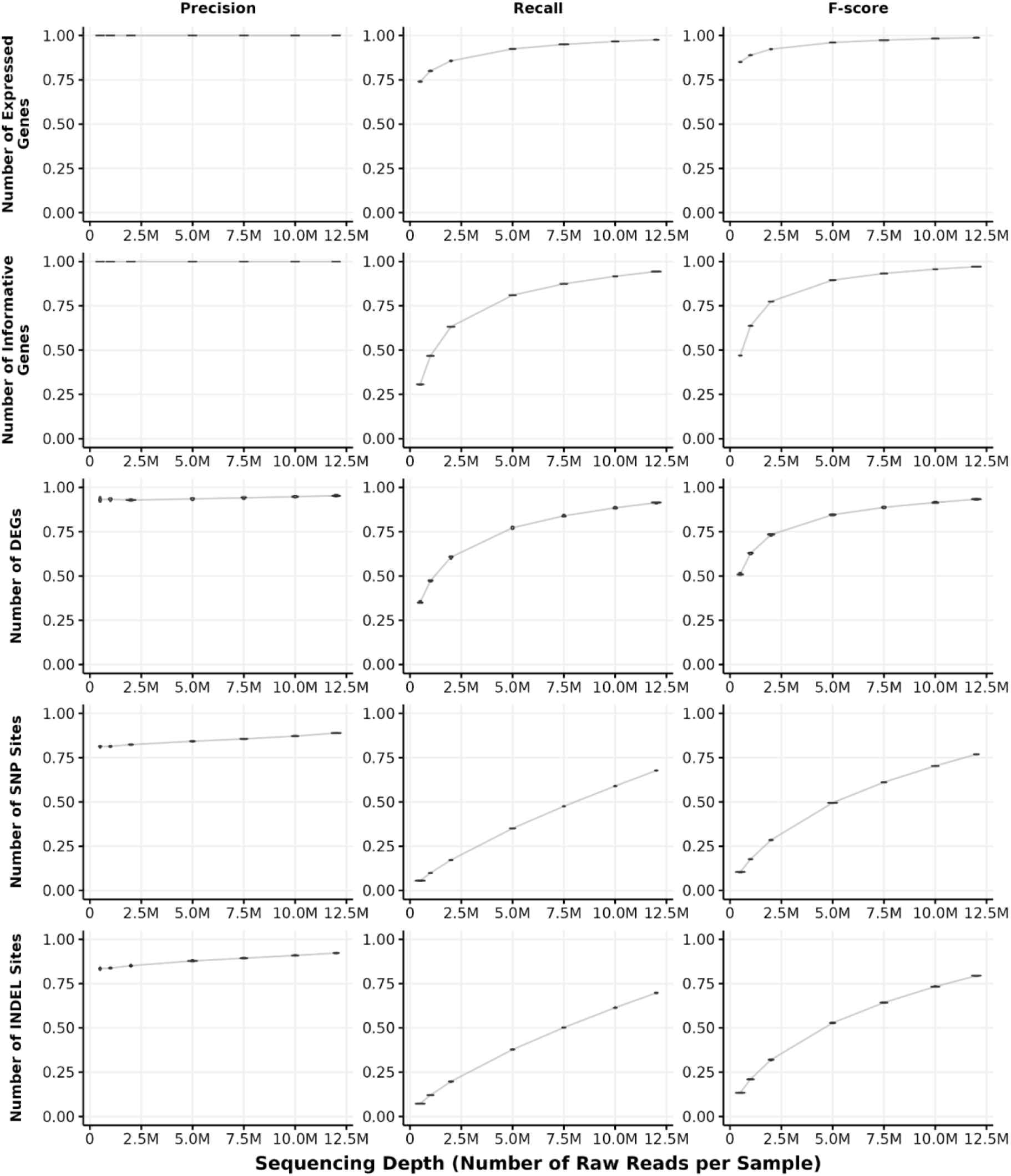
Precision (first column), Recall (second column), and F-score (third column) for expressed genes (first row), informative genes (second row), differentially expressed genes (DEGs)(third row), SNPs (fourth row), and INDELS (fifth row) as a function of sequencing depth (number of reads per sample).

**Figure S4:**
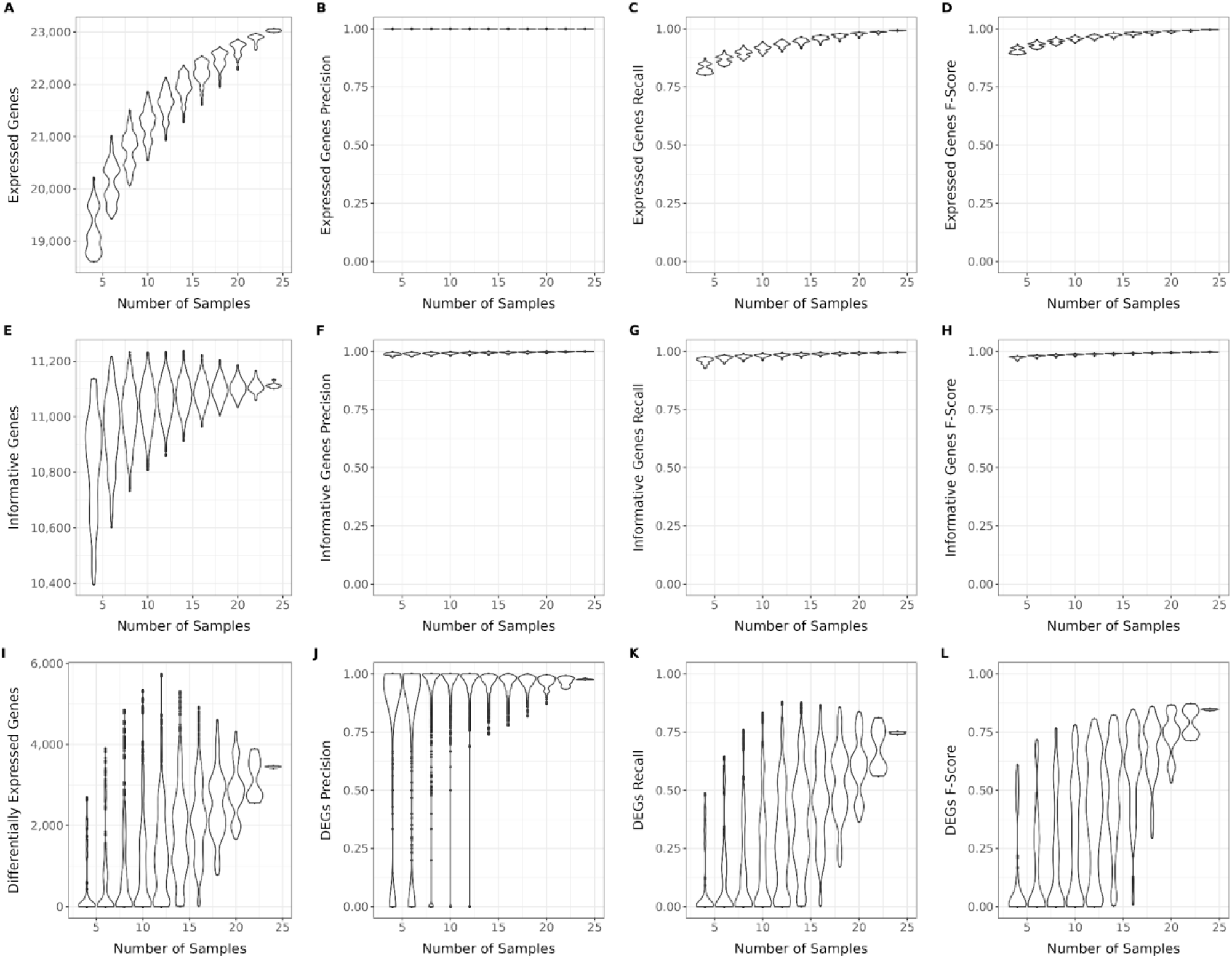
Effect of sample size on the number of expressed genes (A-D), informative genes (E-H), and differentially expressed genes (I-L).

**Table S1:** Study design of the 3′ mRNA-Seq libraries used to sequence 29 biological samples representing 15 calves before and after exposure to heat stress for 12 hours. There are 27 samples sequenced with the Takara library and 27 samples sequences with the Lexogen library.

| Calf | Thermoneutral control | Heat stress |
| --- | --- | --- |
| 1 | Takara & Lexogen | Takara & Lexogen |
| 2 | Takara | Takara & Lexogen |
| 3 | Takara & Lexogen | Takara & Lexogen |
| 4 | Takara & Lexogen | Takara & Lexogen |
| 5 | Takara & Lexogen | Takara |
| 6 | Takara & Lexogen | Takara & Lexogen |
| 7 | Takara & Lexogen | Takara & Lexogen |
| 8 | Takara & Lexogen | Takara & Lexogen |
| 9 | Takara & Lexogen | --- |
| 10 | Lexogen | Takara & Lexogen |
| 11 | Takara & Lexogen | Lexogen |
| 12 | Takara & Lexogen | Takara & Lexogen |
| 13 | Takara & Lexogen | Takara & Lexogen |
| 14 | Takara & Lexogen | Takara & Lexogen |
| 15 | Takara & Lexogen | Takara & Lexogen |

## Notes

### Competing Interest Statement

The authors have declared no competing interest.

